# Adaptive receptor expression and the emergence of disease as loss of signaling homeostasis

**DOI:** 10.64898/2026.03.21.713376

**Authors:** Irina Kareva

## Abstract

Our bodies have evolved to maintain homeostasis through regulatory systems that continuously adapt to keep physiological processes within a normal range. From this perspective, complex chronic disease can be understood as a breakdown of compensatory mechanisms, resulting in loss of homeostasis. Here we propose that adaptive receptor expression dynamics may serve as one such compensatory mechanism, increasing receptor surface expression when external ligand is insufficient, and clearing it when signaling is excessive. To explore this, we adapt a previously published agent-based model and simulate it under a range of scenarios. We find that the system of adaptive receptor expression is robust to oscillatory perturbations but not to chronic stress. We propose that receptor turnover dynamics may be better understood as an adaptive, environmentally responsive process rather than a fixed biological property, and that in some cases, disease manifests only after compensatory mechanisms have been pushed past their limits. We conclude with a discussion of implications for understanding complex chronic diseases, for thinking about epigenetic and mutational change as escalating layers of adaptation, and for how we model receptor dynamics in the context of receptor-mediated drug activity.

## Introduction

Our bodies have evolved to maintain homeostasis through regulatory systems that keep everything in balance. For instance, blood glucose is maintained at approximately 100 mg/dL through a bidirectional feedback mechanism: when blood sugar rises after a meal, the pancreas secrete insulin to facilitate glucose uptake into cells; when it falls, the liver is stimulated to release glucose into circulation through gluconeogenesis (Hatting et al. 2018). Similarly, the immune system has evolved to balance protection against external pathogens and internal threats such as cancer, while simultaneously preventing autoimmunity through a complex, mutually regulated network of feedback loops realized by cells and cytokines (Murphy & Weaver 2016; Dettmer 2021). From this perspective, it is reasonable to hypothesize that disease occurs when something becomes dysregulated, i.e., when there is too much or too little of a protein, a signal, or a process, and the feedback loops that normally enforce homeostasis can no longer keep up.

In drug development, one of the central premises of receptor targeting is that disease reflects a disruption of normal physiological signaling, and that therapeutic intervention aims to restore balance. For instance, in HER2-positive breast cancer, Human Epidermal Growth Factor Receptor 2 (HER2) is overexpressed on the tumor cell surface, driving excess proliferative signaling; trastuzumab targets HER2 directly to reduce that signal and slow tumor growth (Slamon et al. 2001). Similarly, in rheumatoid arthritis, Tumor Necrosis Factor alpha (TNF-α) is chronically elevated, perpetuating joint inflammation; drugs like adalimumab block TNF-α to dampen the excessive inflammatory signal and restore immune balance (Feldmann et al. 1995; Zamora-Atenza et al. 2014). In both cases, the underlying mechanistic hypothesis is the same: identify where the regulatory system has been disrupted and intervene to restore a more physiologically normal signaling state.

One of the most fundamental calculations in dose projections for drug development is receptor occupancy (Durham & Blanco 2015; Simon et al. 2013), i.e., how much drug is needed to engage the target sufficiently to mitigate pathological signaling and restore homeostasis. The required level of occupancy varies considerably by mechanism; in some cases, partial target engagement is sufficient or even intentional, as with beta-blockers in heart failure, where full beta-adrenergic blockade can lead to bradycardia, hypotension and clinical worsening (Packer 1998; Lechat et al. 1998), while in others, such as rituximab targeting CD20 in B cell lymphoma, near-complete occupancy is needed to maximally deplete the malignant B cell population (Mao et al. 2013). The latter reflects what is known as a “no regrets” or maximal pharmacology approach, where we aim to saturate the target for the full dosing interval, so that in the event of no observed efficacy, it is because the mechanism is wrong for this indication, rather than the dose being insufficient (Samineni et al. 2024). This strategy is not ideal and can lead to overexposure and increased safety risk. As a result, the field is increasingly moving toward establishing a more mechanistic occupancy-efficacy relationship early in the drug development process, seeking the minimum effective occupancy rather than the maximum achievable one. Doing this well requires a more sophisticated understanding of both the disease mechanism and the underlying receptor dynamics.

Regardless of which strategy is chosen, if receptor occupancy is part of the dose projection calculation, we need to track not just drug but also target concentrations and how they change over time. Receptors are not static entities; they are dynamic, and their surface expression is continuously shaped by internalization, shedding, and recycling back to the cell surface, among other mechanisms. In most mathematical models of receptor-mediated drug activity, we account for this by assuming some intrinsic turnover rate, i.e., a fixed rate of clearance and resynthesis that maintains a homeostatic baseline level of receptor expression.

There are, however, cases where this assumption visibly breaks down. For example, Epidermal Growth Factor Receptor (EGFR), which is targeted by a monoclonal antibody cetuximab to treat metastatic colorectal and head and neck cancer (Peña-Cabia et al. 2022), undergoes accelerated internalization and degradation upon drug binding (Okada et al. 2017), resulting in surface receptor levels meaningfully lower than would be predicted by the standard baseline turnover model. These dynamics are something that must be explicitly accounted for in PKPD models to make meaningful dose projections (Park et al. 2017; Chung et al. 2021). Such cases are often treated as exceptions requiring special handling in an otherwise standard modeling framework. However, here we propose that this might not be an exception but instead an instance that reveals a more general biological pattern, which is that receptor turnover dynamics are not a fixed biological property but part of a larger adaptive process.

Mechanistically, the most likely explanation is that these dynamics reflect an evolved system for maintaining signaling homeostasis: receptor expression increases when the external signal is too low, and decreases (through internalization, shedding, or other clearance mechanisms) when the external signal is too high. That is, the rates of internalization, shedding, and resurfacing are not genetically pre-determined properties of a receptor but manifestations of a more dynamic mechanism that responds to its environment to maintain homeostasis (Von Zastrow & Sorkin 2021). This type of regulation is known as a negative feedback loop, which, unlike positive feedback loops which amplify perturbations, acts to stabilize signaling in the face of external fluctuations (Modell et al. 2015). From that perspective, one can think of the emergence of disease as occurring when these compensatory mechanisms are no longer keeping up with the external signaling, leading to an inability to maintain homeostasis signaling.

To better understand how such a regulatory system might have evolved, it is useful to look at other biological systems that face the same challenge of maintaining homeostasis in the face of external fluctuations. One compelling example is how honeybee colonies regulate the temperature of their hive. To survive and reproduce, bees must maintain hive temperature within a narrow range. They do this through a remarkably simple but effective behavioral mechanism: when the hive gets too cold, bees huddle together and generate heat; when it gets too warm, they spread out and fan to cool it down (Stabentheiner et al. 2021; Stabentheiner et al. 2010). Each bee has individual temperature thresholds for these behaviors, tied to a genetically linked trait.

Interestingly, hives with greater genetic diversity, and therefore a wider spread of individual thresholds, maintain much more stable temperatures, adjusting to external temperature fluctuations gradually, compared to genetically uniform hives (Miller & Page 2009; Jones et al. 2004). Importantly, however, this system has limits: if the external temperature exceeds what the bees can collectively compensate for (there is a limit to how much they can spread out to cool off or how much they can huddle to generate heat), the hive temperature loses homeostasis, which can lead to collapse of the colony (Chen et al. 2026).

We propose here that perhaps cell receptors may be adapting similar mechanisms in regulating signaling, not through fanning or huddling, but through turnover, shedding and internalization, to maintain normal physiological signaling within a homeostatic range. And just as the robustness of the beehive thermostat depends on the diversity of individual bee thresholds, the robustness of receptor-mediated signaling regulation may similarly depend on the heterogeneity of receptor sensitivities across the cell surface.

Critically, just as the beehive system has a breaking point, so too may this receptor regulatory system: there is likely a range of external fluctuations that the cell can compensate for, beyond which the mechanisms can no longer keep up and signaling falls outside the normal range. It is plausible, within this framework, that this is when we would see the emergence of disease. A generalized schematic of this hypothesized process is shown in Figure 1.

**Figure 1.**
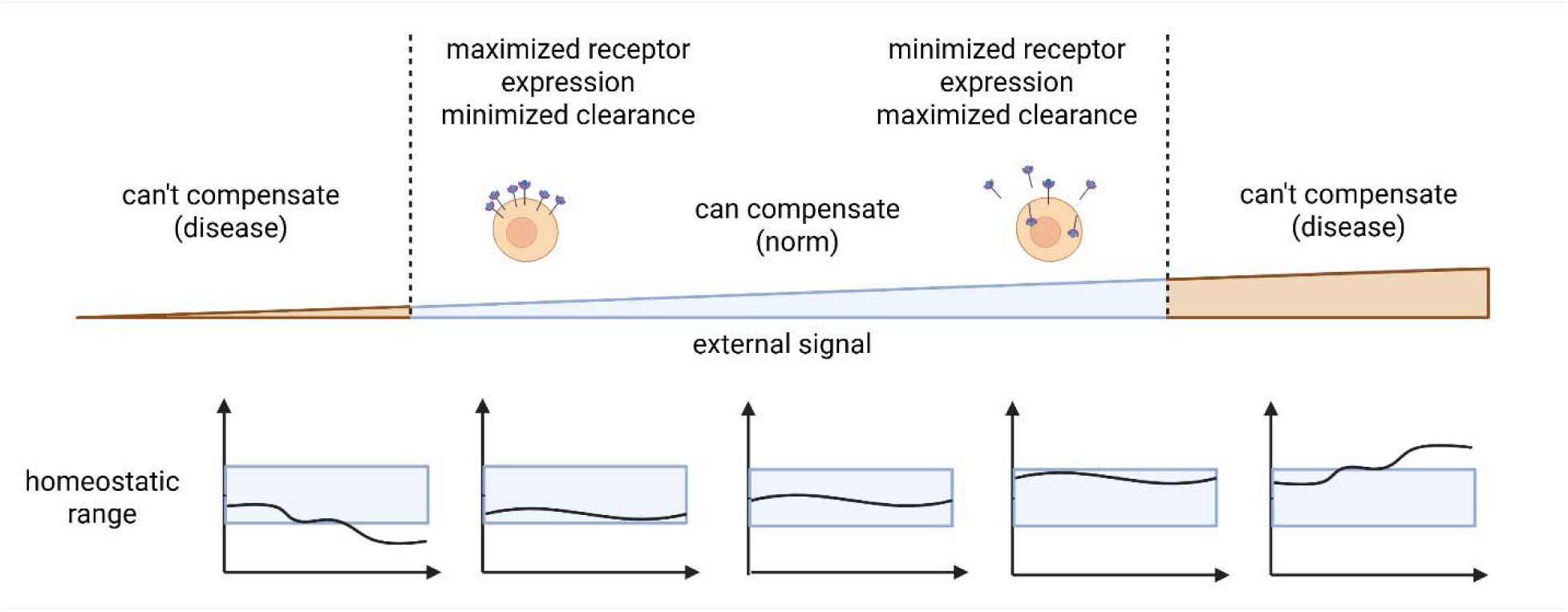
Proposed theory of emergence of complex disease as loss of homeostasis due to inability to compensate for too great a variability in external signaling.

An agent-based model of beehive thermal regulation was previously published in the model library of the agent-based software NetLogo (Wilensky 1999; Page 2025); the original model code was obtained directly from Scott Page (personal communication, 2025). In this manuscript, we first reproduce the original beehive model in MATLAB to establish the baseline system. We then adapt the model structure to describe homeostatic receptor signaling, replacing thermal regulation with receptor expression dynamics, while preserving the underlying logic of distributed threshold responses. We use this adapted model to test the robustness of the system to different types of external perturbations, as well as to explore the impact of interventions. We conclude with a discussion of the implications of these results.

## Methods

### The original beehive thermal regulation agent-based model

First, let us reproduce the original beehive temperature regulation model, originally published in the NetLogo model library (Wilensky 1999; Page 2025) and referenced in (Miller & Page 2009) and recreated here in MATLAB.

The model simulates a bee colony maintaining hive temperature. It is seeded with 120 bees on a spatial grid. Each bee is characterized by two thresholds: a “too warm” threshold, which triggers the bee to spread out and fan to cool the hive, and a “too cold” threshold, which triggers it to cluster and generate heat. Threshold values for each bee are sampled from a normal distribution with a pre-specified mean and variance, giving the population a spread of individual sensitivities.

Notably, the original model includes two subpopulations of bees: younger “ectothermic” bees (55% of the colony), which have limited capacity for metabolic heat generation and fanning, and older “endothermic” bees (45%), which have stronger metabolic heat generation capacity. The older bees are also assumed to have a higher mean “too warm” threshold compared to the younger bees. These assumptions from the original model are preserved here for the purposes of demonstrating that the MATLAB implementation reproduces the original model behavior.

The simulation is conducted as follows. Hive temperature is initialized at 33°C. External temperature varies on a fixed schedule: 20°C → 28°C → 35°C → 10°C → 15°C. At each timestep, if hive temperature falls below a bee’s “too cold” threshold, that bee moves inward and generates heat, slightly raising hive temperature; younger bees contribute less heat than older bees. If hive temperature rises above a bee’s “too warm” threshold, the bee moves outward and fans, slightly lowering hive temperature. The simulation runs for 700 timesteps. Results are shown in Figure 2. The code necessary to reproduce all the simulations can be found at https://github.com/ikareva/beehives.

**Figure 2.**
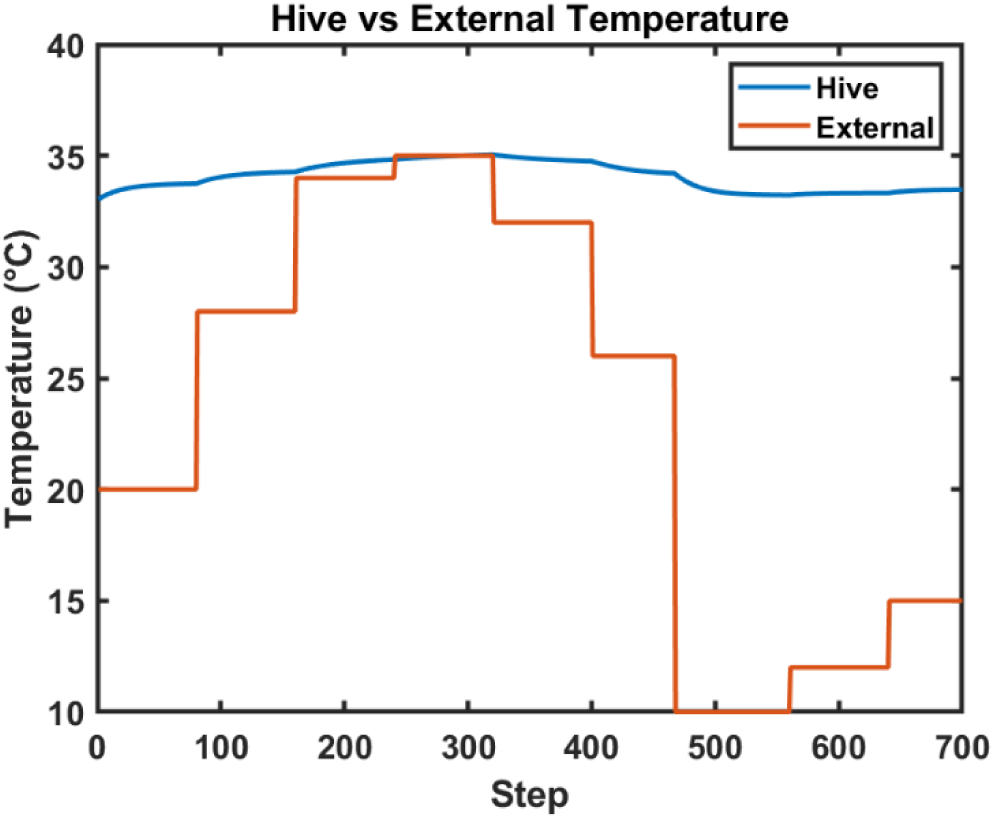
Beehive external vs internal temperature as reproduced by the Matlab version of the bee thermal regulation agent-based model.

### Receptor expression modification of the beehive model

Next, we adapt the beehive thermal regulation model to describe cellular receptor dynamics. The simulation takes place on a grid representing the cell surface, seeded with 120 receptors at fixed spatial positions. In this framework, external ligand concentration corresponds to external temperature in the beehive model, and cell signaling level corresponds to hive temperature. Just as bees responded to hive temperature by adjusting their behavior, receptors respond to cell signaling by modulating their surface expression. A receptor can upregulate (increase surface expression to boost signaling), internalize (decrease surface expression to reduce signaling), or shed (permanently exit the regulatory pool). Both internalization and shedding function as clearance mechanisms that reduce downstream signaling, while upregulation increases it.

Each receptor is characterized by two thresholds: a “too high” threshold, which triggers internalization to reduce signaling output, and a “too low” threshold, which triggers upregulation to increase it. Threshold values are sampled from a normal distribution with a pre-specified mean and variance, giving the receptor population a spread of individual sensitivities. Each receptor also has a baseline expression level (e0).

Analogously to the two subpopulations in the beehive model, receptors are divided into two groups: “baseline” receptors (∼55%), which contribute weakly to signaling regulation, and “regulatory” receptors (∼45%), which have stronger upregulation and internalization capacity. The regulatory receptors are assumed to have a higher mean “too high” threshold compared to baseline receptors.

The full conceptual mapping between the original beehive model structures and the receptor signaling model are summarized in Table 1. Parameter values were chosen arbitrarily for simulation purposes.

**Table 1.** Conceptual mapping of the key concepts between the original beehive thermal regulation model and the adaptive receptor signaling model.

| **Beehive Thermal Regulation Model** | **Homeostatic Receptor Signaling Model** |
| --- | --- |
| External temperature | External ligand concentration |
| Hive temperature | Cell signaling level |
| Bees (agents) | Receptors (agents) |
| Spatial position (radial distance $r$) | Surface expression level ($e \in [0,1]$) |
| Moving inward (clustering) | Upregulating (increasing $e$) |
| Moving outward + fanning (dispersing and active cooling) | Internalizing (decreasing $e$, active signal reduction) |
| Heat generation | Upregulation (signal amplification) |
| Younger bees (ectothermic, 55%) | Baseline receptors (weak regulators, 55%) |
| Older bees (endothermic, 45%) | Regulatory receptors (strong regulators, 45%) |
| "Too cold" threshold | "Too low" threshold |
| "Too warm" threshold | "Too high" threshold |
| Comfortable zone (drift to baseline position) | Comfortable zone (drift to baseline expression $e_0$) |
| N/A (no bee death in original model) | Shedding (permanent receptor removal) |
| Temperature homeostasis target ($\sim 34-36^\circ\text{C}$) | Signaling homeostasis target ($\sim 32-36$ a.u.) |

The simulation is conducted as follows. Cell signaling is initiated at 33 arbitrary units (a.u.). External ligand concentration varies according to one of several predefined scenarios explored in this manuscript, including mild and extreme oscillations, consistently low or high concentrations, chronic increase, and chronic increase with intervention. At each timestep, if cell signaling falls below an individual receptor’s “too low” threshold, that receptor upregulates by increasing its surface expression e, which contributes to raising the signaling level. If cell signaling rises above a receptor’s “too high” threshold, the receptor internalizes by decreasing its surface expression e, which contributes to lowering the signaling level. When signaling is within the comfort band, receptors drift back toward their baseline expression level e0.

The model also includes an optional shedding mechanism, which can be set to one of three modes: disabled (no shedding occurs), conditional (deterministic shedding when a receptor’s expression is near-zero and signaling pressure remains high), or probabilistic (chance-based shedding under similar conditions). Once shed, a receptor is permanently removed from the regulatory pool, with its expression fixed at zero. The simulation runs for 1200 timesteps.

## Results

First, we demonstrate how distributed receptor thresholds maintain cell signaling within a homeostatic range despite external perturbations.

Figure 3A shows the receptor states at the last timepoint of the simulation at t=1200, colored by their regulatory behavior in response to cell signaling levels. Figure 3B shows the external ligand dynamics over time, following the same piecewise schedule used in the original beehive model. Figure 3C tracks how receptor expression patterns respond to these external perturbations. As external ligand increases from t=0 to approximately t=400, the system compensates by reducing upregulation (green band declining), while increasing internalization (orange band rising) and shedding (gray band accumulating). When external ligand drops between approximately t=400 and t=600, the pattern reverses: internalization decreases (orange declining), upregulation increases (green rising), and shedding stabilizes at a low level. As one can see, the external ligand dynamics (Figure 3B) and the corresponding receptor dynamics (Figure 3C) almost mirror each other, highlighting the gradual proportionality of the receptor response. Note also that the final external ligand level is below the original starting level (Figure 3B), and the total receptor expression is proportionally higher than at the beginning of the simulation (black dashed line in Figure 3C), further illustrating how the two systems track each other.

**Figure 3.**
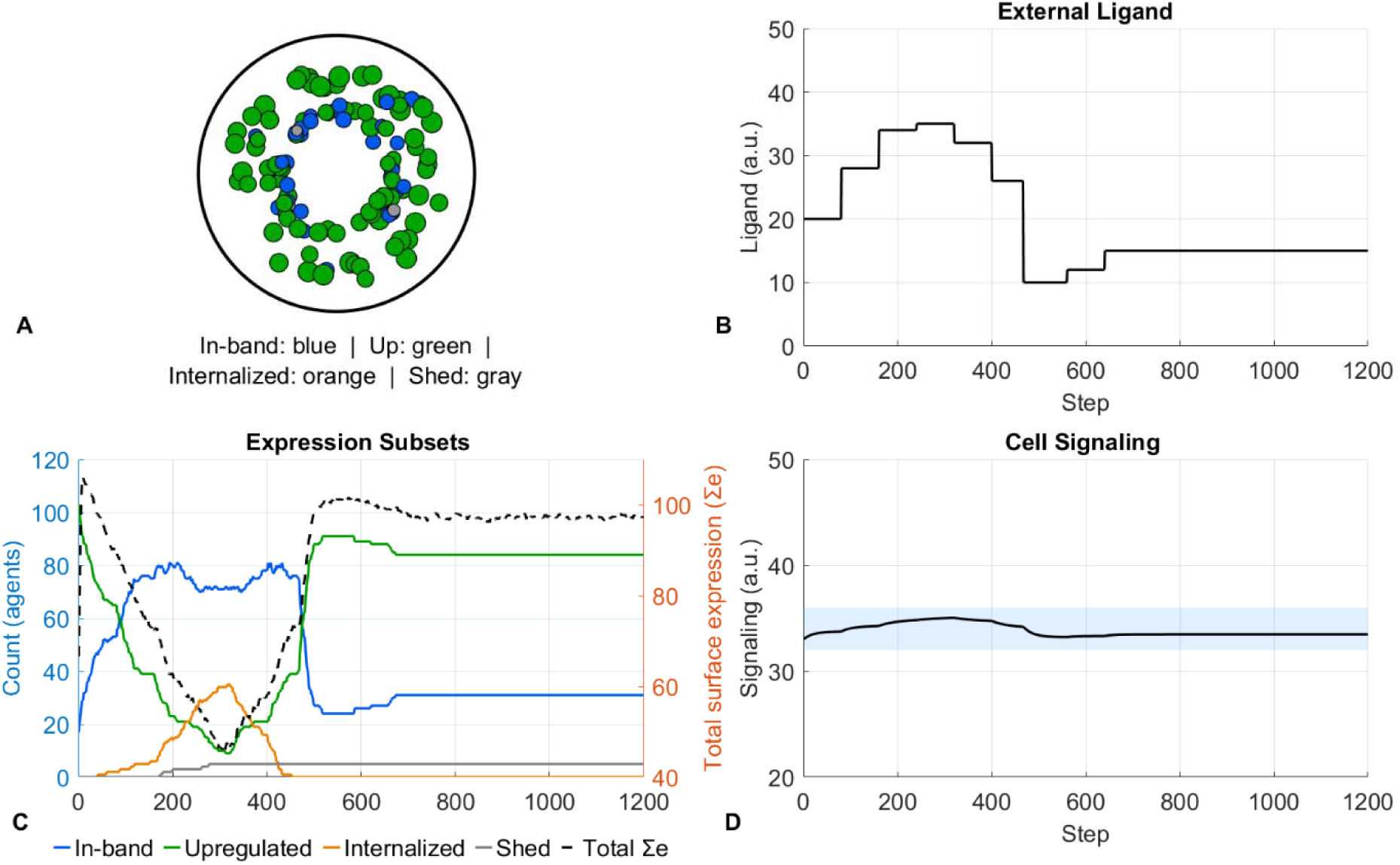
Homeostatic compensation under piecewise external ligand perturbation. (A) Schematic of the cell surface at the end of the simulation. Each circle represents a receptor, colored by regulatory state: blue, within comfort band; green, upregulated; orange, internalized; gray, shed. (B) External ligand concentration over time. This corresponds to external temperature in the original beehive model. (C) Receptor expression dynamics over time. Left y-axis: number of receptors in each regulatory state (colors as in A). Right y-axis: total surface expression (black dashed line), calculated as the sum of individual receptor expression levels. (D) Cell signaling level over time. The light blue band represents the homeostatic signaling range; the black line shows the resulting cell signaling level.

Most importantly, as one can see in Figure 3D, despite large fluctuations in external ligand, cell signaling remains stable within the homeostatic range (light blue band). Just as in the beehive model, the heterogeneity in individual receptor threshold sensitivities allows for a well-regulated, proportional response to external perturbation.

Next, we can use this framework to ask the following question: in a system that has evolved to be robust to perturbations, what does it take to push signaling outside of the normal range? That is, under what conditions would such a system no longer be able to maintain homeostasis?

Notably, we are not considering here the effect of mutations, which would represent a more structural change to the underlying system. Instead, we will explore different external ligand scenarios to identify conditions that can push signaling outside of the homeostatic range through external perturbation alone.

In what follows we will consider six scenarios: mild oscillations; extreme oscillations; consistently low external ligand; consistently high external ligand; chronic increase; and chronic increase with intervention.

### Cases 1 and 2: mild and extreme oscillations

First, let us consider two cases of mild and extreme oscillations in the concentration of the external ligand (Figure 4 and Figure 5). In the mild oscillations scenario, the range of oscillations was set between 26 and 34 arbitrary units (a.u.); in the extreme oscillations scenario, the range is between 18 and 38 a.u.

**Figure 4.**
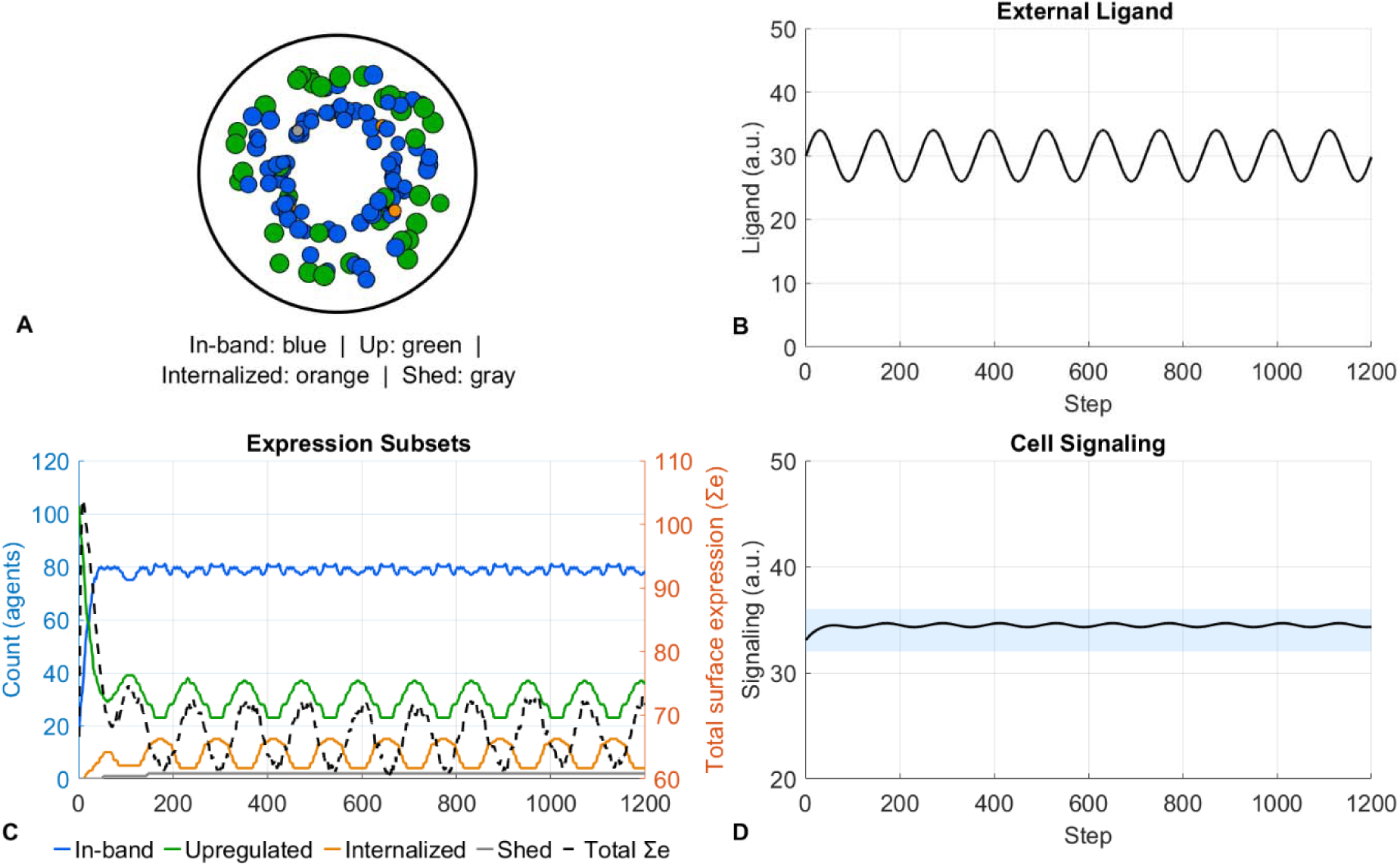
Cell signaling homeostasis under mild external ligand oscillations. (A) Schematic of the cell surface at the end of the simulation. Each circle represents a receptor, colored by regulatory state: blue, within comfort band; green, upregulated; orange, internalized; gray, shed. (B) External ligand concentration over time, oscillating between 26 and 34 a.u. (C) Receptor expression dynamics over time. Left y-axis: number of receptors in each regulatory state (colors as in A). Right y-axis: total surface expression (black dashed line), calculated as the sum of individual receptor expression levels. (D) Cell signaling level over time. The light blue band represents the homeostatic signaling range; the black line shows the resulting cell signaling level.

**Figure 5.**
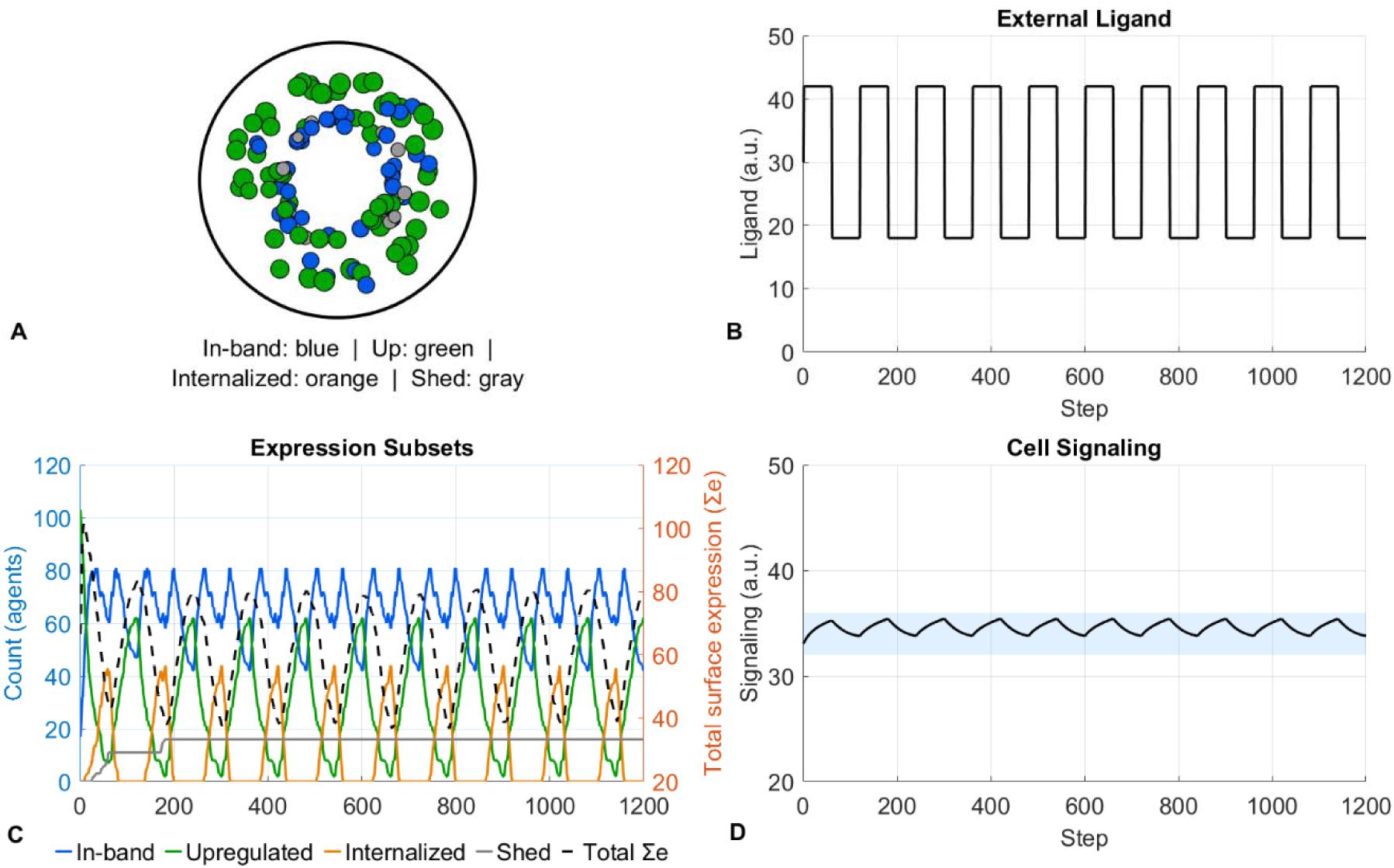
Cell signaling homeostasis under extreme external ligand oscillations. (A) Schematic of the cell surface at the end of the simulation. Each circle represents a receptor, colored by regulatory state: blue, within comfort band; green, upregulated; orange, internalized; gray, shed. (B) External ligand concentration over time, oscillating between 18 and 38 a.u. (C) Receptor expression dynamics over time. Left y-axis: number of receptors in each regulatory state (colors as in A). Right y-axis: total surface expression (black dashed line), calculated as the sum of individual receptor expression levels. (D) Cell signaling level over time. The light blue band represents the homeostatic signaling range; the black line shows the resulting cell signaling level.

In the first scenario, as one can see, most receptors remain within their stable range (blue, Figure 4A). The patterns of receptor upregulation and clearance (Figure 4C) predictably mirror the pattern of external ligand dynamics (Figure 4B), and downstream cell signaling remains stably within the normal band (Figure 4D).

In the second scenario, where the amplitude of oscillations is much larger (Figure 5B), the compensatory mechanisms are considerably more dynamic (Figure 5C).

Interestingly, total receptor expression is on average higher and has a higher amplitude in the extreme oscillations case compared to the mild one (Figure 4C vs Figure 5C), even though downstream signaling remains firmly within the normal range in both cases (Figure 4D vs Figure 5D).

Together, these two cases highlight that while signaling remains within the homeostatic range in both cases regardless of oscillation amplitude, the upstream receptor dynamics required to achieve that outcome can look quite different. Importantly, despite the oscillatory behavior of the external ligand, the system proves robust in both cases.

### Case 3. Consistently low external ligand concentrations

Next, let us consider a scenario where the concentration of the external ligand is consistently low. The goal of this analysis is to see whether, within this system, one could push signaling below the normal range by dramatically reducing ligand concentration.

External ligand concentration was set between 2 and 6 a.u. As one can see in Figure 6, the system compensated by maximally upregulating receptor expression (green, Figure 6A and Figure 6C). Despite the external ligand concentration being substantially lower than in the previous cases (average of 4 a.u. vs approximately 30 a.u.), downstream signaling remained within the normal range, albeit at the lower end (Figure 6D). Notably, total receptor expression in this scenario is at its highest relative to all previous cases (black dashed line, Figure 6C), reflecting the system doing everything it can to amplify the available signal by maximizing receptor surface expression to “squeeze out” as much downstream signaling as possible from very little external ligand.

**Figure 6.**
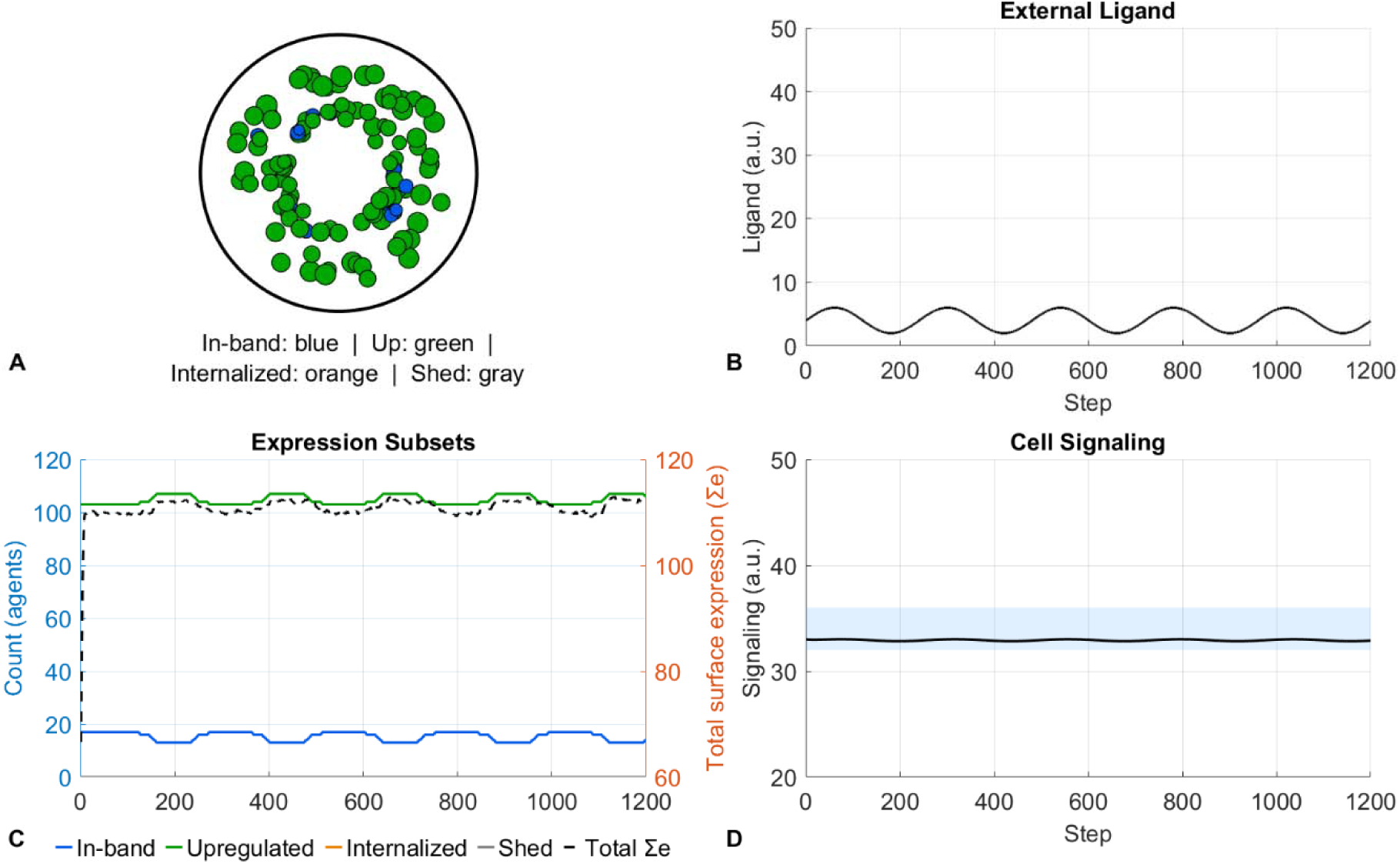
Cell signaling response to consistently low external ligand concentrations. (A) Schematic of the cell surface at the end of the simulation. Each circle represents a receptor, colored by regulatory state: blue, within comfort band; green, upregulated; orange, internalized; gray, shed. (B) External ligand concentration over time, maintained between 2 and 6 a.u. (C) Receptor expression dynamics over time. Left y-axis: number of receptors in each regulatory state (colors as in A). Right y-axis: total surface expression (black dashed line), calculated as the sum of individual receptor expression levels. (D) Cell signaling level over time. The light blue band represents the homeostatic signaling range; the black line shows the resulting cell signaling level.

While this is of course a single parametrization, it nevertheless highlights that the compensatory mechanisms of receptor expression regulation can be remarkably robust to external variations, at least on the lower bound.

### Case 4. Consistently high external ligand concentrations

Now let us evaluate a scenario when the concentration of the external ligand is consistently high. The concentration of the external ligand is taken to be between 40 and 44 a.u.

In this scenario, we begin to observe the system’s compensatory mechanisms finally failing to keep up. As one can see in Figure 7A and C, receptor upregulation has been fully suppressed (green band tending to zero), while the clearance mechanisms are maximally engaged (orange internalization and gray shedding at their highest levels). There are still some receptors for whom this level of signaling remains within their individual range (blue), but total receptor expression is substantially lower compared to the previous case (black dashed line, Figure 6C vs Figure 7C). Notably, this is the first scenario in which downstream signaling rises outside of the normal range (Figure 7D), indicating that, just as in the beehive where consistently high external temperatures eventually exceed the bees’ capacity to cool the hive, here the receptors cannot be downregulated any further to keep signaling within the homeostatic range.

**Figure 7.**
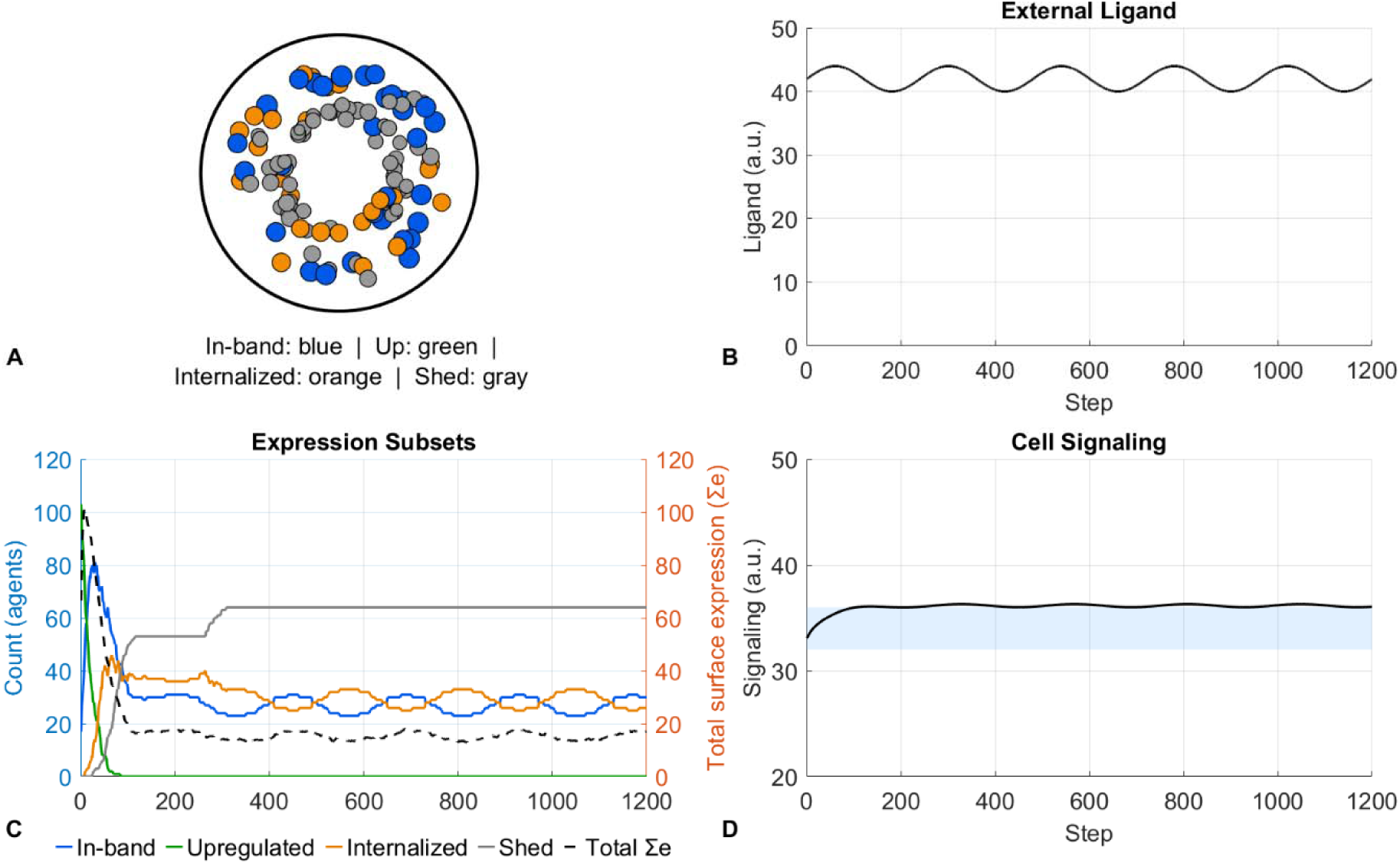
Cell signaling response to consistently high external ligand concentrations. (A) Schematic of the cell surface at the end of the simulation. Each circle represents a receptor, colored by regulatory state: blue, within comfort band; green, upregulated; orange, internalized; gray, shed. (B) External ligand concentration over time, maintained between 40 and 44 a.u. (C) Receptor expression dynamics over time. Left y-axis: number of receptors in each regulatory state (colors as in A). Right y-axis: total surface expression (black dashed line), calculated as the sum of individual receptor expression levels. (D) Cell signaling level over time. The light blue band represents the homeostatic signaling range; the black line shows the resulting cell signaling level.

Next let is consider an extension of this example, when the external ligand is chronically increasing.

### Cases 5 and 6. Chronic increase with and without intervention

Now let us consider the last two cases, depicting the impact of a chronically increasing external ligand on the system, and the effect of a transient intervention on that same system. The external ligand concentration starts at 15 a.u. and increases to stabilize at 44 a.u.; as everywhere, these values were chosen arbitrarily for illustrative purposes.

The first case is shown in Figure 8. This is similar to the consistently high scenario (Case 4), except here we can see a more gradual picture of how the system adapts to a continuous increase in external ligand concentration. As one can see in Figure 8C, the system first responds by suppressing upregulation (green falling), followed by progressive increases in internalization (orange) and shedding (gray). Total receptor expression stabilizes at a low level relative to the beginning of the simulation (black dashed line), and downstream signaling eventually rises outside of the normal range (Figure 8D). Arguably, this can be interpreted as the emergence of disease, where abnormal signaling arises not from mutations or any structural change to the system, but purely from compensatory mechanisms no longer being able to keep up with the changes in the external environment.

**Figure 8.**
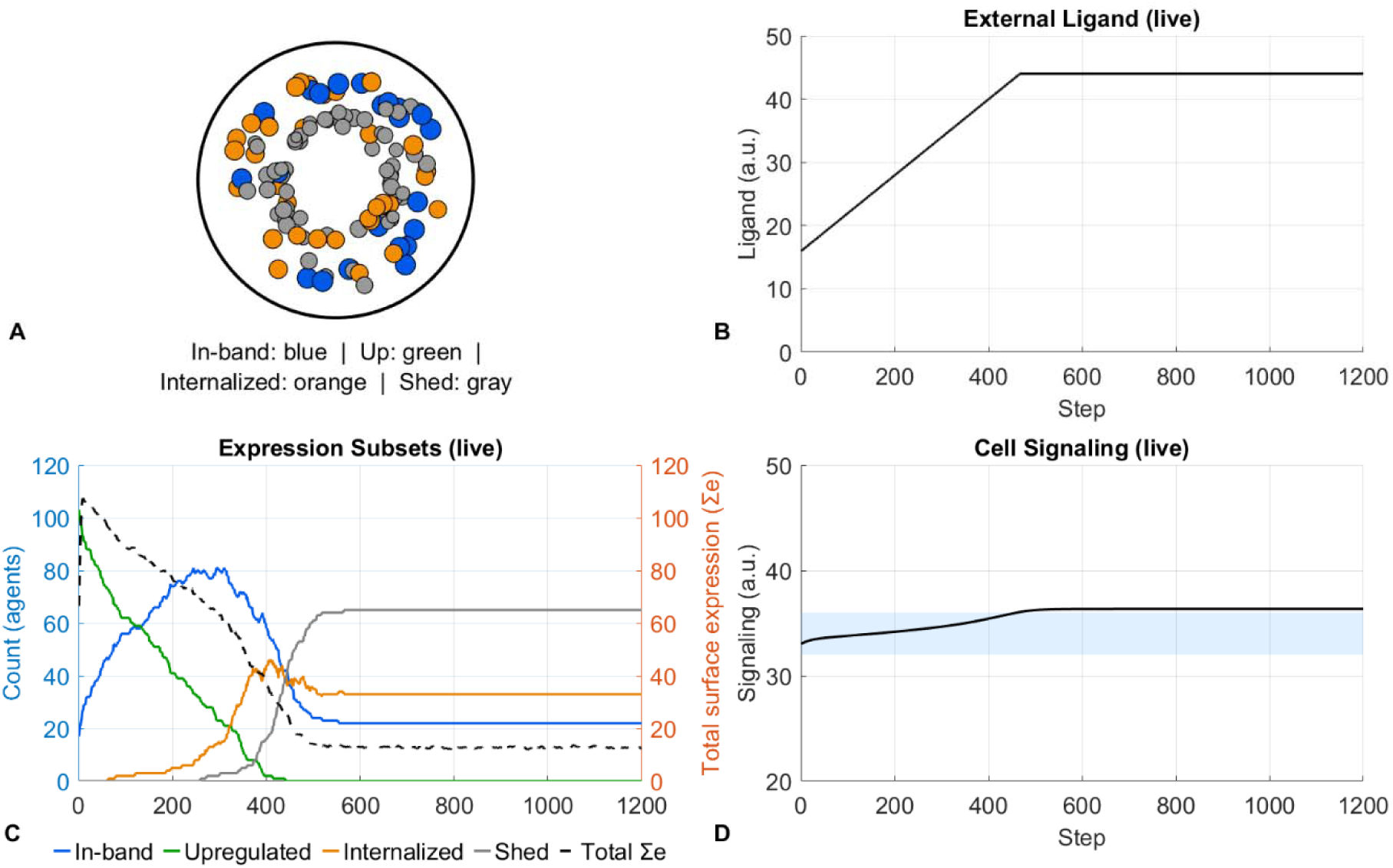
Cell signaling response to chronic external ligand increase. (A) Schematic of the cell surface at the end of the simulation. Each circle represents a receptor, colored by regulatory state: blue, within comfort band; green, upregulated; orange, internalized; gray, shed. (B) External ligand concentration over time, increasing from 15 to 44 a.u. (C) Receptor expression dynamics over time. Left y-axis: number of receptors in each regulatory state (colors as in A). Right y-axis: total surface expression (black dashed line), calculated as the sum of individual receptor expression levels. (D) Cell signaling level over time. The light blue band represents the homeostatic signaling range; the black line shows the resulting cell signaling level.

Now let us explore a variation of this scenario, where some of the ligand is removed from the external ligand pool at an arbitrarily chosen timepoint t=320, mimicking a therapeutic intervention that clears the ligand.

As one can see in Figure 9, the receptor expression system temporarily recalibrates after the intervention; however, without any mechanism sustaining the reduced ligand concentration, the system reverts to high levels of abnormal signaling. The intervention does delay the onset of abnormal signaling, occurring at approximately t=800 with intervention compared to approximately t=500 without it; however, without addressing the underlying causes of the continuous ligand increase, this arguably mimics treating the symptom rather than the cause.

**Figure 9.**
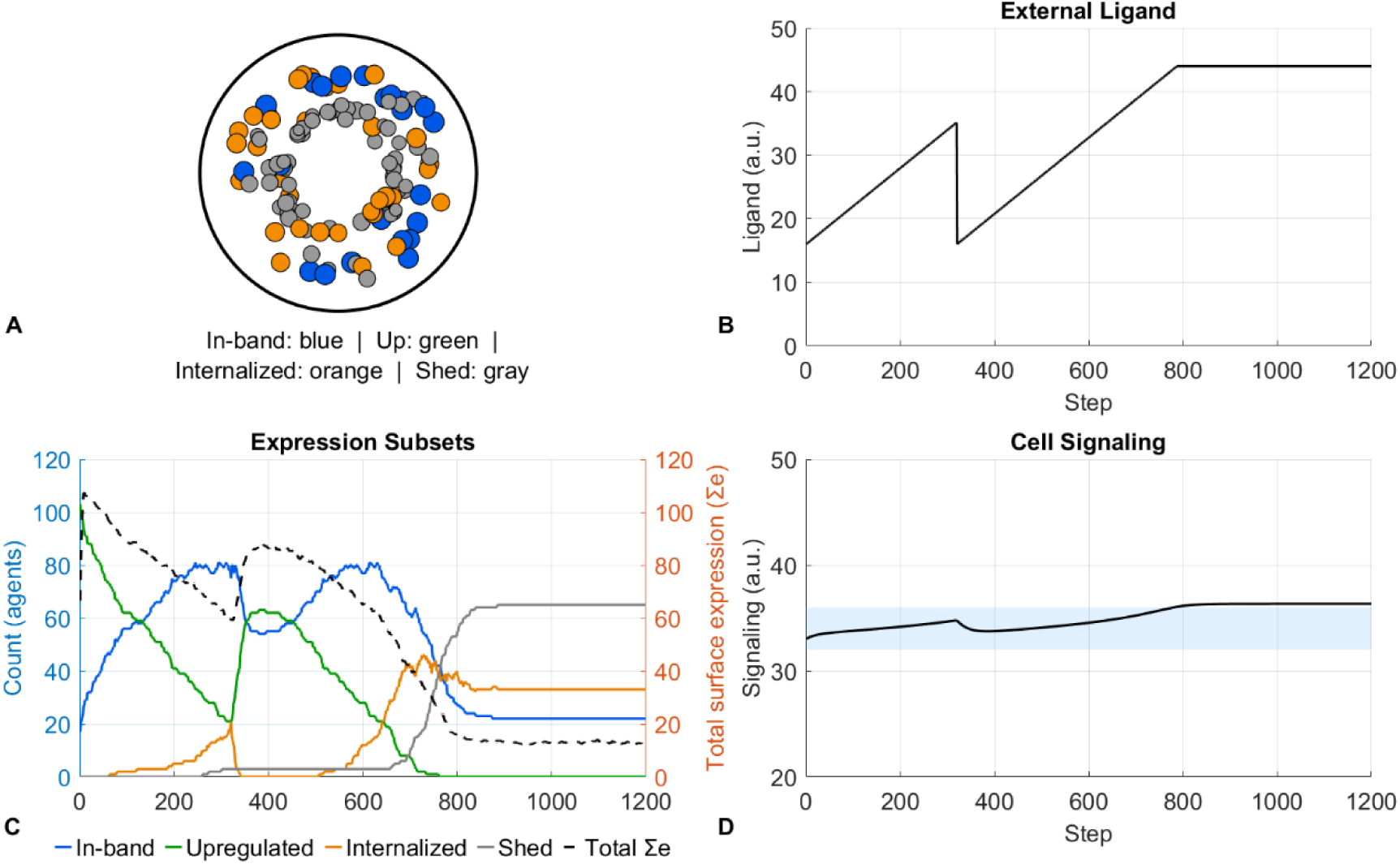
Cell signaling response to chronic external ligand increase with therapeutic intervention. (A) Schematic of the cell surface at the end of the simulation. Each circle represents a receptor, colored by regulatory state: blue, within comfort band; green, upregulated; orange, internalized; gray, shed. (B) External ligand concentration over time, increasing from 15 to 44 a.u., with a transient reduction at t=320 mimicking ligand clearance by therapeutic intervention. (C) Receptor expression dynamics over time. Left y-axis: number of receptors in each regulatory state (colors as in A). Right y-axis: total surface expression (black dashed line), calculated as the sum of individual receptor expression levels. (D) Cell signaling level over time. The light blue band represents the homeostatic signaling range; the black line shows the resulting cell signaling level.

## Discussion

In this paper we explored the concept of homeostasis and how adaptive receptor expression dynamics may be involved in maintaining it, with the goal of understanding complex chronic disease as a disruption of homeostatic signaling. We built upon an analogy between beehive thermal regulation and cellular receptor dynamics: just as bees huddle when the hive gets too cold and spread out when it gets too warm, we proposed that receptors on the cell surface upregulate when external ligand is too low and clear when it is too high. We adapted a previously published agent-based beehive thermal regulation model to this framework and tested the robustness of such a system to a range of external ligand scenarios, interpreting loss of signaling homeostasis as emergence of disease.

We found that such a system is remarkably hard to push out of homeostasis. It proved robust to a wide range of perturbations and across a wide magnitude of fluctuations.

Particularly striking was the lower bound, where even when external ligand concentration was reduced to very low levels, the system was able to compensate by maximizing receptor expression and minimizing all clearance mechanisms, still keeping the downstream signal within the normal range, albeit near its lowest bound. Pushing the system out of homeostasis on the upper bound, however, was achievable through chronically elevated or continuously increasing external ligand. Once all clearance mechanisms are exhausted, i.e., when the receptors are maximally shed or internalized, and when receptor upregulation is maximally suppressed, the signal can rise outside of the normal range.

We also simulated a transient intervention, physically removing some ligand from the system at a single timepoint. Predictably, the system temporarily recalibrated and signaling returned toward the homeostatic range; however, since the underlying driver of ligand increase remained ongoing, the system eventually reverted to abnormal signaling. This is analogous to treating the symptom rather than the underlying cause. This could be analogous to a therapy that requires lifelong administration simply to keep signaling within the normal range.

One compelling analogy to this framework can be found in the emergence of type 2 diabetes (T2D). When food is ingested, blood glucose rises, prompting the pancreas to secrete insulin, which binds to the insulin receptor on the cell surface and causes translocation of the GLUT4 glucose transporter, allowing glucose to enter the cell and be stored, maintaining blood glucose at approximately 100 mg/dL (Stuart et al. 2009); this is not the only glucose clearance-mechanism but for the purposes of this discussion we will focus on insulin-mediated glucose clearance. When the amount of glucose entering the system exceeds the body’s storage capacity, the pancreas compensates by producing more insulin for longer. As was shown in the landmark work of Kraft (Kraft & others 1975), subsequently re-analyzed on a larger dataset by Crofts et al. (Crofts et al. 2016), hyperinsulinemia precedes hyperglycemia: the body works increasingly hard to keep blood glucose within the normal range, and it is only when these compensatory mechanisms can no longer keep up that a person transitions from normal glucose tolerance through pre-diabetes to overt T2D. Critically, as was argued in (Kareva 2024), this process can be understood not just as a primary failure of the insulin-receptor signaling machinery but additionally as a consequence of diminishing capacity to store excess glucose, particularly in skeletal muscle, the largest and most modifiable storage depot. Within this framework, the pancreas is doing exactly what our receptor model does under chronically elevated external ligand, which is compensating maximally until it cannot anymore.

The therapeutic implications follow the same logic as well. Interventions that assist the failing compensatory mechanisms, such as injecting exogenous insulin, stimulating the pancreas with sulfonylureas, or clearing glucose directly from the blood with SGLT-2 inhibitors (Del Prato & Pulizzi 2006; Chao & Henry 2010), can keep signaling within the normal range, but without addressing the underlying cause of chronic glucose excess, they are treating the symptom rather than the root problem. This distinction has consequences beyond diabetes itself: chronic hyperinsulinemia, which can be present even in individuals with normal fasting glucose as shown by Crofts et al. (Crofts et al. 2016), has been shown to act as a growth signal for cancer cells, and Hopkins et al. (Hopkins et al. 2018) demonstrated that a drug-induced diabetic phenotype, and specifically the resulting hyperinsulinemia, caused resistance to PI3K inhibitors in mouse tumor models, which was reversed upon normalization of insulin levels. This suggests that the downstream consequences of a compensatory system pushed past its limits extend well beyond the primary disease, and that addressing the underlying dysregulation rather than its symptoms may have broader therapeutic implications than currently appreciated.

The framework proposed here that in some cases disease emerges when compensatory mechanisms can no longer maintain homeostatic signaling in the face of chronic external perturbation, may extend well beyond receptor dynamics and offer a useful lens for thinking about complex chronic diseases more broadly. The T2D analogy discussed above is one instance of this pattern. In oncology, a structurally analogous process may underlie the emergence of T cell exhaustion: as we explored in (Kareva & Gevertz 2024), both underlying inflammation and possibly chronic antigen stimulation from a growing tumor can drive progressive transitions from functional cytotoxic T cells through reversibly exhausted states to terminally exhausted ones. The immune system is compensating, adapting, trying to maintain a balance between cytotoxic activity and autoimmune protection until the chronic inflammatory signal, augmented by the therapy, overwhelms those mechanisms, and it is mitigating this additional inflammation that can help the immune system recalibrate, restoring treatment efficacy (Mathew et al. 2024; Zak et al. 2024). In Alzheimer’s disease, a similar argument can be made. As reviewed in (Kareva 2025), amyloid plaques and tau tangles may be better understood not as the root cause of neurodegeneration but as late-stage byproducts of an underlying metabolic collapse. That is, one can see these adaptations as the brain’s compensatory mechanisms for managing energy deficit and neuroinflammation, which eventually fail after years of chronic increase of external stress. In both of these cases, the disease does not appear suddenly but instead it emerges after the compensatory systems have been working at or near their limits for a long time. This suggests that there may be a window, during which early warning signals are present but the system has not yet broken down, just as hive temperature can serve as an early warning signal of potential upcoming honeybee colony collapse (Colin et al. 2025). Identifying and acting on those signals, rather than waiting for overt disease, may be one of the most important implications of this framework.

Interestingly, within this framework we were able to observe adaptive behavior and emergence of disease without invoking any assumptions about mutations. Instead, here disease (as interpreted through the lens of loss of homeostasis) emerges purely from compensatory mechanisms no longer being able to keep up with chronic external perturbation.

This raises an intriguing evolutionary hypothesis. One could think of the body’s response to chronic environmental challenges as operating in layers of escalating commitment. The first layer is the one we have modeled here, which is through fast and reversible compensatory mechanisms, such as receptor upregulation and clearance, which continuously adjust to maintain signaling homeostasis. If these mechanisms are sufficient, homeostasis is preserved and no lasting change occurs. If they are not sufficient, one could hypothesize that the next layer of adaptation is epigenetic, where more durable changes in gene expression do not alter the underlying DNA sequence but reconfigure the cell’s regulatory landscape in response to the persistent signal.

Stress-induced epigenetic remodeling is well-documented, and the idea that chronic signaling drives epigenetic adaptation is consistent with emerging evidence across multiple biological contexts, from metabolic to neurodevelopmental to mental health (Dion et al. 2022; Kubota 2016; Faraji & Metz 2026).

Further, it’s possible that if even epigenetic adaptation proves insufficient and the chronic perturbation continues, one could speculate that this creates conditions that increase the likelihood of mutational change. It’s possible that this could be a mechanism to trigger not random mutation in the classical sense, but mutation nudged by the failure of every preceding compensatory layer to restore homeostasis. The idea that cellular stress can promote mutagenesis is not without precedent. For instance, McClintock’s classic observations on transposable elements activating under stress (McClintock 1984; Maggert 2019), and more recent work on error-prone DNA repair under chronic inflammatory conditions (Kay et al. 2019), suggest that the genome itself may be more plastic under stress than the standard model of random mutation implies. A schematic representation of this hypothesis is shown in Figure 10.

**Figure 10.**
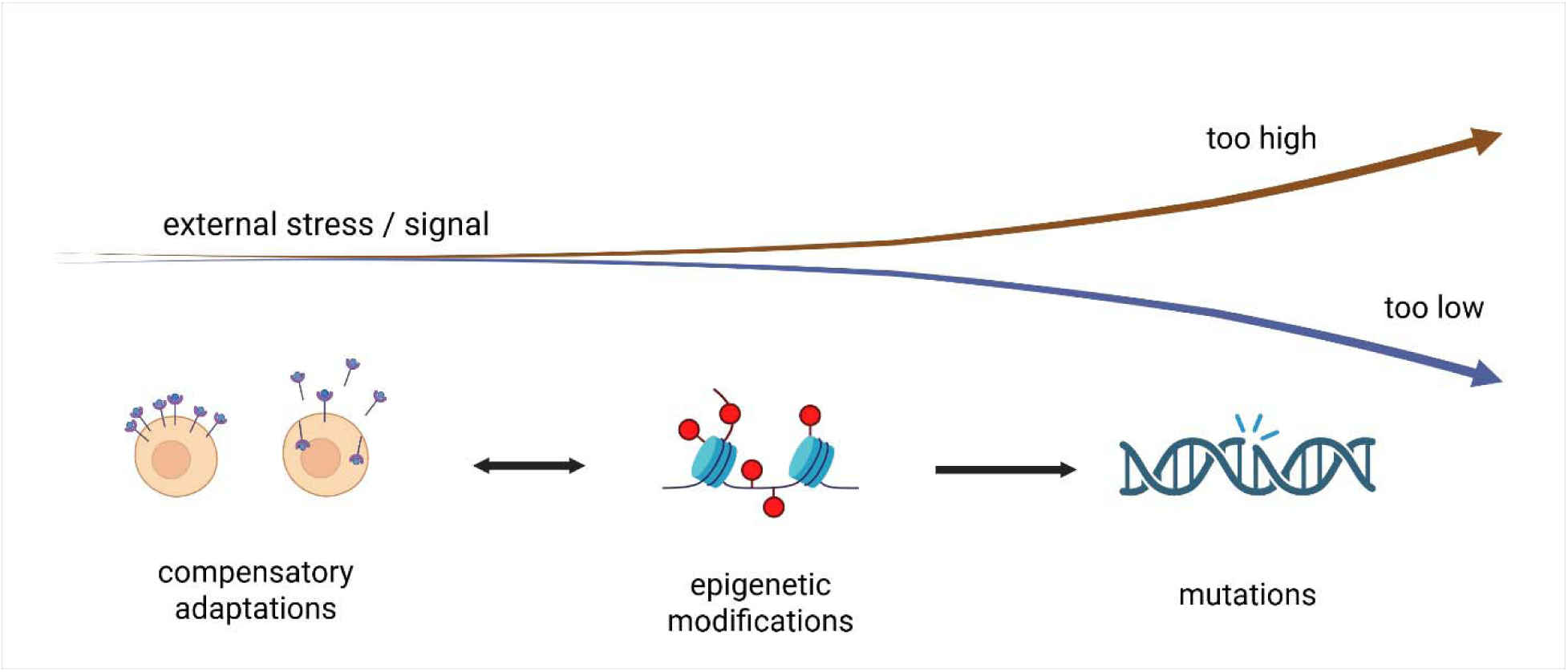
Hypothesis that compensatory mechanisms are the first layer of response to chronic stress.

Within this framework, one could think of mutation not only as a cause of disease but also as a last-resort adaptive response. This reframing has potentially important implications. It suggests that the window for intervention may be longer than we think, and that preventing the exhaustion of earlier compensatory layers, rather than waiting for mutations to appear, could be a more effective therapeutic strategy.

This hypothesis is, of course, speculative, but it is testable. A conceptually clean experiment would be to expose a cell population to chronically elevated ligand concentrations (mimicking the conditions of Cases 4 and 5 in our model) and to then longitudinally track both epigenetic marks, such as DNA methylation patterns and mutation rates, as compensatory mechanisms, which are expected to be progressively engaged and then exhausted. If the hypothesis holds, one would expect to see epigenetic drift preceding any mutational changes, with the degree of drift correlating with the degree of signaling dysregulation. Such an experiment would not only test the proposed hierarchy of adaptation but could also help identify early epigenetic signatures that precede overt disease.

As with any mathematical model, this work has important limitations that warrant acknowledgment. The model is not calibrated to specific experimental data or a particular molecular mechanism; parameters were chosen for illustrative purposes, and the simulation is intended to explore qualitative behavior rather than make quantitative predictions. There is no mechanistic explanation embedded in the model for why a receptor would internalize under some conditions and shed under others, nor for what drives the specific patterns of external ligand variation we simulated. The model does capture heterogeneous receptor contributions through two discrete subpopulations (baseline receptors with weaker regulatory capacity and regulatory receptors with stronger upregulation and internalization responses) but this is a simplification of what in reality is likely a continuous distribution of receptor sensitivities. Furthermore, the model treats signaling as a lumped global variable, and does not capture spatial organization of receptors, clustering effects (Garcia-Parajo & Mayor 2024), co-receptor interactions (Mørch et al. 2020), or downstream pathway saturation (Altszyler et al. 2014), all of which are known to play important roles in receptor-mediated signaling in real biological systems. These are meaningful simplifications, and calibrating this framework to specific receptor systems and experimental data would be an important next step.

Nevertheless, even as a conceptual framework, this model raises questions that we think are worth taking seriously. Chief among them is whether our standard approach to mathematical modeling of receptor dynamics in drug development needs revisiting. We typically assume that receptors have some fixed homeostatic expression level and some intrinsic turnover rate, and we model drug binding against that static backdrop.

But if receptor expression is itself an adaptive variable that responds to the signaling environment rather than simply maintaining a predetermined level, then our models may be systematically misrepresenting the system we are trying to describe.

This reframing has practical and well-supported implications for understanding drug resistance. Compensatory receptor upregulation in response to targeted therapy has been documented across multiple oncology settings. For instance, HER3 upregulation has been observed in response to HER2 blockade (Li et al. 2018), MET amplification compensates for EGFR inhibition in lung cancer (Engelman et al. 2007), and FGFR upregulation occurs in response to VEGFR inhibition (Ichikawa et al. 2020). Additionally, androgen receptor amplification and upregulation is a well-characterized mechanism of resistance to androgen deprivation therapy in prostate cancer (Takeda et al. 2018), which is a clear example of the prediction made here, that when you suppress the signal, the system responds by upregulating the receptor.

If these are not exceptions but manifestations of a general adaptive principle, then the clear next question is whether one could therapeutically exploit such adaptation. For instance, could a transient suppression of a growth factor signal, which may be insufficient on its own for tumor control, be used to drive compensatory receptor upregulation, and then a receptor-targeting agent such as an antibody-drug conjugate (ADC) administered precisely when receptor expression is at its peak, thereby maximizing target availability and potentially the therapeutic effect? A possible experiment to test this hypothesis would be to treat a tumor model with a short course of low-dose ligand-blocking therapy, monitor surface receptor expression levels over time using flow cytometry or imaging, identify the timepoint of maximal receptor upregulation, and administer a receptor-targeting agent at that moment compared to standard timing. If the hypothesis holds, one would expect to see improved efficacy from the sequenced approach, using the cell’s compensatory response to effectively sensitize a tumor to be more receptive to therapy. This is a complementary approach to the one suggested by (Gatenby et al. 2025), where the authors used natural selection to resensitize prostate cancer tumors to androgen deprivation therapy.

More broadly, these questions point toward a shift in how we might think about drug targets, away from treating disease as a collection of dysregulated levers to be individually corrected, and toward asking why those levers evolved to be there in the first place, and what happens when we pull them without understanding the system they are part of. We hope that this work, and the questions it raises, will contribute to that conversation.

## Acknowledgements

The author would like to thank Dr. Scott Page, from whose lectures on “Understanding Complexity” she learned (among many other things) about mechanisms of beehive thermal regulation, and who in generously shared his copy of the beehive thermal regulation code upon request in October 2025. This work has not received any external funding. The views expressed in this work are the author’s own and do not represent the views of any specific organization or funding agency.

## Code availability

Matlab code necessary to reproduce all the results reported in this manuscript is available in https://github.com/ikareva/beehives.

## Use of AI

ChatGPT was used to convert to Matlab the original code shared in October 2025 by Scott Page of the original beehive thermal regulation model. All the outputs were manually verified by IK.

## References

Altszyler, E. et al., 2014. Impact of upstream and downstream constraints on a signaling module’s ultrasensitivity. Physical biology, 11(6), p.066003.

Chao, E.C. & Henry, R.R., 2010. SGLT2 inhibition—a novel strategy for diabetes treatment. Nature reviews drug discovery, 9(7), pp.551–559.

Chen, J. et al., 2026. Negative Effects of Excessive Heat on Colony Thermoregulation and Population Dynamics in Honeybees. Ecological and Evolutionary Physiology, 99(1), pp.000–000.

Chung, T.K. et al., 2021. A target-mediated drug disposition population pharmacokinetic model of GC1118, a novel anti-EGFR antibody, in patients with solid tumors. Clinical and Translational Science, 14(3), pp.990–1001.

Colin, T. et al., 2025. Temperature as an early warning signal of honeybee colony failure. Ecological Informatics, p.103445.

Crofts, C. et al., 2016. Identifying hyperinsulinaemia in the absence of impaired glucose tolerance: An examination of the Kraft database. Diabetes research and clinical practice, 118, pp.50–57.

Dettmer, P., 2021. Immune: A journey into the mysterious system that keeps you alive, Random House.

Dion, A., Munoz, P.T. & Franklin, T.B., 2022. Epigenetic mechanisms impacted by chronic stress across the rodent lifespan. Neurobiology of stress, 17, p.100434.

Durham, T.B. & Blanco, M.-J., 2015. Target engagement in lead generation. Bioorganic & medicinal chemistry letters, 25(5), pp.998–1008.

Engelman, J.A. et al., 2007. MET amplification leads to gefitinib resistance in lung cancer by activating ERBB3 signaling. science, 316(5827), pp.1039–1043.

Faraji, J. & Metz, G.A., 2026. Environmental epigenetics: new horizons in redefining biological and health outcomes. Environment International, p.110072.

Feldmann, M. et al., 1995. TNF alpha is an effective therapeutic target for rheumatoid arthritis. Annals of the New York Academy of Sciences, 766, pp.272–278.

Garcia-Parajo, M.F. & Mayor, S., 2024. The ubiquitous nanocluster: A molecular scale organizing principle that governs cellular information flow. Current opinion in cell biology, 86, p.102285.

Gatenby, R.A. et al., 2025. Directed Evolution Restored Castrate Sensitivity in a Patient With Castrate Resistant Metastatic Prostate Cancer. The Prostate, 85(16), pp.1562–1567.

Hatting, M. et al., 2018. Insulin regulation of gluconeogenesis. Annals of the New York Academy of Sciences, 1411(1), pp.21–35.

Hopkins, B.D. et al., 2018. Suppression of insulin feedback enhances the efficacy of PI3K inhibitors. Nature, 560(7719), pp.499–503.

Ichikawa, K. et al., 2020. Activated FGF2 signaling pathway in tumor vasculature is essential for acquired resistance to anti-VEGF therapy. Scientific reports, 10(1), p.2939.

Jones, J.C. et al., 2004. Honey bee nest thermoregulation: diversity promotes stability. Science, 305(5682), pp.402–404.

Kareva, I., 2024. From hyperinsulinemia to cancer progression: how diminishing glucose storage capacity fuels insulin resistance. Aging and Cancer, 5(3), pp.51–61.

Kareva, I., 2025. The many roads to dementia: a systems view of Alzheimer’s disease. arXiv preprint arXiv:2504.05441.

Kareva, I. & Gevertz, J.L., 2024. Mitigating non-genetic resistance to checkpoint inhibition based on multiple states of immune exhaustion. npj Systems Biology and Applications, 10(1), p.14.

Kay, J. et al., 2019. Inflammation-induced DNA damage, mutations and cancer. DNA repair, 83, p.102673.

Kraft, J.R. & others, 1975. Detection of diabetes mellitus in situ (occult diabetes). Lab med, 6(2), pp.10–22.

Kubota, T., 2016. Epigenetic alterations induced by environmental stress associated with metabolic and neurodevelopmental disorders. Environmental Epigenetics, 2(3), p.dvw017.

Lechat, P. et al., 1998. Clinical effects of β-adrenergic blockade in chronic heart failure: a meta-analysis of double-blind, placebo-controlled, randomized trials. Circulation, 98(12), pp.1184–1191.

Li, X. et al., 2018. Posttranscriptional upregulation of HER3 by HER2 mRNA induces trastuzumab resistance in breast cancer. Molecular Cancer, 17(1), p.113.

Maggert, K.A., 2019. Stress: An evolutionary mutagen. Proceedings of the National Academy of Sciences, 116(36), pp.17616–17618.

Mao, C.-P. et al., 2013. Subcutaneous versus intravenous administration of rituximab: pharmacokinetics, CD20 target coverage and B-cell depletion in cynomolgus monkeys. PLoS One, 8(11), p.e80533.

Mathew, D. et al., 2024. Combined JAK inhibition and PD-1 immunotherapy for non–small cell lung cancer patients. Science, 384(6702), p.eadf1329.

McClintock, B., 1984. The significance of responses of the genome to challenge. Science, 226(4676), pp.792–801.

Miller, J.H. & Page, S.E., 2009. Complex adaptive systems: an introduction to computational models of social life: an introduction to computational models of social life, Princeton university press.

Modell, H. et al., 2015. A physiologist’s view of homeostasis. Advances in physiology education.

Mørch, A.M. et al., 2020. Coreceptors and TCR signaling–the strong and the weak of it. Frontiers in cell and developmental biology, 8, p.597627.

Murphy, K. & Weaver, C., 2016. Janeway’s immunobiology, Garland Science.

Okada, Y. et al., 2017. EGFR downregulation after anti-EGFR therapy predicts the antitumor effect in colorectal cancer. Molecular Cancer Research, 15(10), pp.1445–1454.

Packer, M., 1998. β-Adrenergic blockade in chronic heart failure: principles, progress, and practice. Progress in cardiovascular diseases, 41(1), pp.39–52.

Page, S.E., 2025. Complex Adaptive Systems (NetLogo model list). Available at: https://sites.lsa.umich.edu/scottepage/home/complex-adaptive-systems/.

Park, W.-S. et al., 2017. Use of a Target-Mediated Drug Disposition Model to Predict the Human Pharmacokinetics and Target Occupancy of GC 1118, an Anti-epidermal Growth Factor Receptor Antibody. Basic & clinical pharmacology & toxicology, 120(3), pp.243–249.

Peña-Cabia, S. et al., 2022. Assessment of exposure-response relationship for cetuximab in patients with metastatic colorectal cancer and head and neck cancer. Farmacia Hospitalaria, 46(1), pp.21–26.

Del Prato, S. & Pulizzi, N., 2006. The place of sulfonylureas in the therapy for type 2 diabetes mellitus. Metabolism, 55, pp.S20–S27.

Samineni, D. et al., 2024. Dose optimization in oncology drug development: an international consortium for innovation and quality in pharmaceutical development white paper. Clinical Pharmacology & Therapeutics, 116(3), pp.531–545.

Simon, G.M., Niphakis, M.J. & Cravatt, B.F., 2013. Determining target engagement in living systems. Nature chemical biology, 9(4), pp.200–205.

Slamon, D.J. et al., 2001. Use of chemotherapy plus a monoclonal antibody against HER2 for metastatic breast cancer that overexpresses HER2. New England journal of medicine, 344(11), pp.783–792.

Stabentheiner, A. et al., 2021. Coping with the cold and fighting the heat: thermal homeostasis of a superorganism, the honeybee colony. Journal of Comparative Physiology A, 207(3), pp.337–351.

Stabentheiner, A., Kovac, H. & Brodschneider, R., 2010. Honeybee colony thermoregulation–regulatory mechanisms and contribution of individuals in dependence on age, location and thermal stress. PLoS one, 5(1), p.e8967.

Stuart, C.A. et al., 2009. Insulin-stimulated translocation of glucose transporter (=) 12 parallels that of = 4 in normal muscle. The Journal of Clinical Endocrinology & Metabolism, 94(9), pp.3535–3542.

Takeda, D.Y. et al., 2018. A somatically acquired enhancer of the androgen receptor is a noncoding driver in advanced prostate cancer. Cell, 174(2), pp.422–432.

Wilensky, U., 1999. NetLogo. Available at: http://ccl.northwestern.edu/netlogo/.

Zak, J. et al., 2024. JAK inhibition enhances checkpoint blockade immunotherapy in patients with Hodgkin lymphoma. Science, 384(6702), p.eade8520.

Zamora-Atenza, C. et al., 2014. Adalimumab regulates intracellular TNFα production in patients with rheumatoid arthritis. Arthritis research & therapy, 16(4), p.R153.

Von Zastrow, M. & Sorkin, A., 2021. Mechanisms for regulating and organizing receptor signaling by endocytosis. Annual Review of Biochemistry, 90(1), pp.709–737.

